# Exploring the impact of intraspecific interaction between sibling colonies of *Bacillus cereus* MSM-S1 in the absence and presence of antibiotics

**DOI:** 10.64898/2026.10.19.760111

**Authors:** Kritika Prasad, Brinta Chakraborty, Saurav K Saha, Tapas K Sengupta

**Affiliations:** Department of Biological Sciences, Indian Institute of Science Education and Research Kolkata, Mohanpur, Nadia - 741 246, West Bengal, India

**Author notes:** Correspondence and reprints”.

**Keywords:** *Bacillus cereus*, Sibling colonies, Quorum sensing, Sporulation, NheA toxin

## Abstract

*Bacillus cereus* is a spore-forming pathogen and food contaminant that can survive in hostile environments through different survival strategies. As part of survival, *B. cereus* may also engage in intra- or interspecific interactions that shape its community structure and stress responses, enabling its persistence under unfavourable conditions. The present study explored intraspecific interactions, at the cell and molecular levels, between two sibling colonies of *Bacillus cereus* MSM-S1 grown through swarming adjacently on nutrient agar in the absence and presence of gentamicin. While growing alone, gentamicin altered the symmetrical colony morphology of *B. cereus* MSM-S1, which was associated with cell elongation, decreased flagellation and increased expression of the toxin gene *nheA*. During interaction, both in the absence and presence of gentamicin, the motility of sibling colonies towards each other was restricted, forming a distinct line/zone of demarcation between the colonies. Microscopic analysis at different time points (day 3, 5 and 7) revealed progressive changes in the swirling arrangement of cells at the interaction zone. In the absence of gentamicin, cells became jammed at the interacting front by day 7, whereas this effect was not observed in the presence of gentamicin on day 7. Cells at the interacting zone underwent gradual sporulation; notably, sporulation was less in the presence of gentamicin. Further analysis suggested that nutrient limitation was the major contributory factor for the formation of the zone of no growth between the interacting colonies. Moreover, the expression of quorum-sensing genes was down-regulated in cells at the interacting zone without gentamicin, but notably, upregulated in the presence of gentamicin. The expression of chemotaxis regulator gene, cheY, was also upregulated in interacting zone cells both in the absence and presence of gentamicin. In addition, increased expression of the global transcriptional regulator codY in the interacting zone under both conditions suggested its involvement in metabolic regulation during sibling colony interaction. The findings highlight modulated quorum sensing and sporulation as adaptive mechanisms in *Bacillus cereus* MSM-S1 to cope with external stress imposed by nutrient limitations caused by the growth of sibling colonies, whereas, cell elongation and toxin production together is a preferred survival strategy for a growing MSM-S1 colony when stress is imposed by gentamicin as antibiotic but sufficient nutrient is available.

## Introduction

*Bacillus cereus*, a spore-forming bacterium, is widespread in the environment and is known to contaminate food. **(Li et al., 2025)**. It is implicated in intestinal and non-intestinal diseases owing to its enterotoxins and tissue-destructive enzymes **(Bottone, 2010)**. Besides food poisoning, it can cause illnesses like endophthalmitis, pneumonia, and gas gangrene-like skin infections in immunocompetent and immunocompromised people **(Bottone, 2010).**

Swarming motility is a social behaviour observed in bacteria in which flagellated bacteria move cooperatively across semi-solid surfaces **(Kearns, 2010)**. Swarming bacteria have been shown to exhibit adaptive or non-genetic resistance to antibiotics compared to planktonic counterparts **(Bhattacharyya et al., 2020; Butler et al., 2010).** Thus, swarming represents a multicellular adaptive strategy in bacteria to overcome environmental stress.

Bacteria are social organisms that respond to external stimuli from their surroundings. Bacterial cells in nature rarely exist solely; instead, they form dense communities that can be composed of single or multiple species **(Davey & O’toole, 2000)**. A wide variety of interactions can occur within microbial communities, including intraspecific interactions (within individuals of the same species) and interspecific interactions (between individuals of different species). Moreover, microbial interactions also influence antibiotic activity and contribute to the emergence of antibiotic resistance **(Muzafar et al., 2026; Zhang et al., 2026)**. Therefore, bacterial interactions can not only influence the growth of microorganisms but also reshape the selective pressures imposed by antibiotics. Hence, the social behaviour of microorganisms is also an important aspect to consider to reveal insights about alterations in motility, such as swarming, stress tolerance, antibiotic resistance and pathogenicity that would be unnoticed by studying bacteria in isolation.

Although interspecific interaction among bacteria has been commonly observed, intriguingly, a rare intraspecific sibling rivalry is also noted among a few bacterial species. For instance, *Paenibacillus dendritiformis*, a Gram-positive bacterium, kills the cells of its neighbouring sibling colonies by secreting inhibitory compounds that become lethal above a threshold **(Be’er et al., 2009).** Sublitisin, an extracellular protease secreted by *P. dendritiformis,* was found to be responsible for the appearance of the inhibition zone between the colonies. In another study, the sibling colonies of *Marinobacter* sp. have also been shown to exhibit a form of non-lethal chemical communication that prevents colonies from overrunning each other’s niche space. A protein, glycerophosphoryl diester phosphodiesterase, was enriched in the inhibition zone between the spreading colonies, which was speculated to be involved in the overall response **(Cruz-López et al., 2021).**

Besides the presence of competing species, the presence of antibiotics in the environment also poses a threat to bacterial survival and impacts the structure and functioning of microbial communities. Antibiotics can have both direct (short-term) and indirect (long-term) effects on microbial communities. The short-term effects include bactericidal and bacteriostatic effects, which may cause some microbial populations and their ecological processes to vanish. The emergence of antibiotic-resistant bacteria and, in certain situations, bacterial strains capable of breaking them down through metabolic or co-metabolic processes are examples of the indirect long-term effects **(Grenni et al., 2018)**.

*B. cereus* often exists not as isolated cells but as spatially structured colonies that can interact with its neighbours, including bacteria from different or the same genera, or even same species, including genetically related species (sibling colonies). The intraspecific sibling interactions may critically influence colony morphology, resource partitioning, competitive dynamics, and collective behaviours like swarming. Also, in response to environmental changes or stress, *B. cereus* modifies its physiology to optimize resource utilization while maintaining structural and genetic integrity by minimizing damage to cellular architecture **(Duport et al., 2016)**.

Thus, the present study attempts to decipher the dynamics of intraspecific interaction between sibling colonies of the isolated *Bacillus cereus* MSM-S1 strain in the absence and presence of gentamicin at the colony, cellular and molecular levels. The strain was isolated from soil but harbours genes for virulence factors and thus can behave as an opportunistic pathogen and presents a good model organism for the study. Overall, the study represents how bacteria can modify their behaviour according to their environment and attempts to unravel the mechanisms employed by bacteria to survive under stress conditions.

## 2. Materials and Methods

### 2.1 Bacterial Strains and Growth Conditions

The isolated strain *Bacillus cereus* MSM-S1 was previously identified by 16S rDNA sequencing (Genbank accession no. HM061612) and deposited in the National Collection of Industrial Microorganisms, CSIR-NCL Pune, India (Strain Deposit Number #NCIM 1212) **(Chakraborty et al., 2016)**. The strains *Pseudomonas aeruginosa* KPW.1-S1 (MTCC10087) **(Dasgupta et al., 2013)** and *Bacillus thuringiensis* KPWP1 (Genbank accession no. FJ 897722) used in the study were isolated and reported previously **(Banerjee et al., 2013)**. All strains were stored and maintained in glycerol stocks (15% glycerol).

For the present study, the strain was grown in Nutrient broth (0.5% Peptone, 0.3% Beef extract, 0.5% NaCl). For swarming and interaction studies, nutrient agar with 0.6% agar (w/v) was used. Gentamicin at the required concentration was added to the media before pouring plates.

### 2.2 Antibiotic susceptibility test and determination of Minimum Inhibitory Concentration of gentamicin

The antibiotic susceptibility test was done by the standard disk diffusion method on nutrient agar medium with 1.5% agar (w/v). For that, *Bacillus cereus* MSM-S1 was grown in nutrient broth medium at 37°C till turbidity developed. 100 μL of bacteria were first spread on nutrient agar plates with 1.5% agar to form a uniform lawn, and an antibiotic disc (Octodisc, HiMedia OD014R-1PK G-IV-minus) was placed on the plates aseptically. Eight different antibiotics were tested, including ampicillin (A, 10μg), cephalothin (Ch, 5 μg), colistin methanesulphonate (Cl, 25 μg), gentamicin (G, 10 μg), streptomycin (S, 10 μg), sulphtriad (Sl, 200 μg), tetracycline (T, 25 μg), and cotrimoxazole (Co, 25 μg). The plates were incubated at 30°C overnight. The result was interpreted as resistant or sensitive based on the appearance of the zone of inhibition surrounding the disc.

The minimum inhibitory concentration of gentamicin was determined following the protocol described by **(Kumar et al., 2023)**. 2mL of *Bacillus cereus* MSM-S1 culture was taken in test tubes with different concentrations of gentamicin sulfate (0.25, 0.5, 0.75, 1, 1.25, 1.5, 2, 2.5, 5, and 10 µg/mL) and incubated at 150 rpm, 37°C for 18 hours. The next day, the O.D. of the cultures was measured at 600nm in a spectrophotometer (Shimadzu UV-1800).

### 2.3 Effect of subinhibitory concentrations of gentamicin on swarming motility of Bacillus cereus MSM-S1

The effect of subinhibitory concentrations of gentamicin on the swarming ability of *Bacillus cereus* MSM-S1 was first determined by spotting 2μL (10^7^ CFU/mL) of *Bacillus cereus* MSM-S1 culture on nutrient agar plates with 0.6% agar (w/v) in the absence and presence of 0.25µg/mL (MIC/40) and 0.5µg/mL (MIC/20) of gentamicin. The bacterial spots were allowed to dry for 15 minutes. The plates were then incubated at 30°C for several days. Images of growing colonies were taken every day in a gel documentation system (SYNGENE G: BOX). The ImageJ version 1.54g was used to calculate the area covered by the growing colonies on each day.

### 2.4 Studies of Interaction between sibling colonies of Bacillus cereus MSM-S1

Studies of the interaction between sibling colonies of *Bacillus cereus* MSM-S1 were performed on nutrient agar plates with 0.6% agar, which allowed for the swarming of bacterial colonies and facilitated interaction between the sibling colonies. For studying the interaction between the sibling colonies in the presence of antibiotics, the plates were prepared with the addition of 0.5μg/mL gentamicin. 2μL (10^7^ CFU/mL) of *Bacillus cereus* MSM-S1 culture was spotted adjacent to each other at a distance of 3cm. Single spots of *Bacillus cereus* MSM-S1 on separate plates were used as a control. The bacterial spots were allowed to dry for 15 minutes and incubated at 30°C for several days. Images of growing colonies were captured daily using a gel documentation system (SYNGENE G: BOX). The ImageJ version 1.54g was used to calculate the area covered by the growing colonies on each day.

### 2.5 Structured Illumination Microscopy using Axio Observer microscope with the ApoTome 2 module

Structured Illumination Microscopy was performed to observe the in situ arrangement of peripheral cells of *Bacillus cereus* MSM-S1 during sibling colony interaction. The edge of the colonies with agar base was cut into equal-sized cubes using a sterile scalpel. Further, the cells in the colonies were stained with 0.01% acridine orange (w/v) and washed twice with 1X PBS. The stained samples were then put on coverglass bottom dishes and were visualized using an Axio Observer microscope with the ApoTome 2 module. The image acquisition was completed with the help of Zen Pro 2012 software, Carl Zeiss (Germany). The samples were excited at 488 nm using an argon laser, and emission of 493 to 576 nm was detected **(Chakraborty et al., 2016)**.

### 2.6 Scanning electron microscopy

The surface topography of *Bacillus cereus* MSM-S1 cells during sibling colony interaction was observed through Field Emission Scanning Electron Microscopy (FESEM). Colonies of *Bacillus cereus* MSM-S1 were grown alone and adjacent to each other at a distance of 3 cm on semi-solid nutrient agar plates with 0.6% agar in the absence and presence of gentamicin. Peripheral cells from the growing colonies in the absence and presence of gentamicin were collected and smeared on round coverslips. 50µL of 2.5% glutaraldehyde (made in PBS) was added to them, followed by incubation for 2 hours at 4°C. The samples were then washed three times with 1X PBS. 50µL of 0.1% osmium tetroxide (made in PBS) was then added to the samples, and then the samples were incubated in the dark for 40 minutes. Samples were then washed three times with 1x PBS. The cells were dehydrated serially by adding 50µL of 30%, 40%, 50%, 60%, 70%, 80%, and 90% ethanol. The samples were kept for 5 minutes each time alcohol was added. Finally, 50µL of 100% ethanol was added and kept for 15 minutes **(Bhadani et al., 2025)**. The excess ethanol was removed by gentle pipetting. Next, the coverslips were air-dried and further allowed to dry in a desiccator. The coverslips containing the samples were coated with a thin layer of conducting metal, specifically gold-palladium, in a sputter coater. Finally, cells were observed under a Supra 55 VP field emission scanning electron microscope (Carl Zeiss), with SmartSEM software, and images were taken **(Chakraborty et al., 2016).**

### 2.7 Transmission Electron Microscopy

To check any alteration in the flagellation of the *Bacillus cereus* MSM-S1 cells during swarming in the absence and presence of gentamicin, transmission electron microscopy (TEM) was performed. For this, bacterial cells were taken from the peripheries of colonies growing in the absence and presence of gentamicin and resuspended in 1X phosphate-buffered saline. 2μL of bacterial suspension was placed on the Formvar-coated copper grid (300 mesh size) and fixed using 2.0% paraformaldehyde. After 15 minutes, excess liquid was wicked off carefully. The grids were stained with 2.0% (w/v) phosphotungstic acid and kept in the dark for 5 minutes. The grids were then air-dried and further dried in a desiccator. Fixed cells were viewed under a transmission electron microscope (JEOL, JEM-2100 plus electron microscope) **(Bhadani et al., 2025)**.

### 2.8 Differential expression of genes

To understand the expression of different genes during intraspecific interaction, sibling colonies of *Bacillus cereus* MSM-S1 were grown adjacent to each other at a distance of 3cm for 7 days in the absence and presence of gentamicin. Peripheral cells of the colony growing alone, from non-interacting and interacting zones of the sibling colonies, were scraped and resuspended in 1XPBS, followed by centrifuging at 5000 rpm for 5 minutes to obtain a cell pellet. Total RNA was isolated from the samples using RNAiso Plus (Takara, Japan) following the protocol mentioned by the manufacturer. The RNA samples by treating them with RNase-free DNase I (Thermo Fisher Scientific, USA) to remove residual genomic DNA. Then, the RNA samples were used for cDNA synthesis using the PrimeScript 1st strand cDNA synthesis kit (Takara, Japan) following the instructions provided by the manufacturer. Briefly, a 10μL reaction mixture was prepared with 1μg of RNA, 2μL of 50 μM Random 6-mers, and 1μL of 10 mM dNTP mixture and was incubated at 65°C for 5 minutes and immediately kept on ice. 4μL of 5X PrimeScript buffer, 0.5μL of 40 U/μL RNase inhibitor, and 1 μL of 200 U/μL PrimeScript reverse transcriptase were added to the initial 10 μL reaction mixture, and the final volume was made up to 20 μL with sterile nuclease-free water. The reaction mixtures were first incubated at 30°C for 10 minutes, followed by incubation at 42°C for 60 minutes, and finally at 70°C for 15 minutes in a Veriti 96-well thermal cycler (Applied Biosystems, Thermo Fisher Scientific, USA). The resulting products were kept at −20°C until use. Real-time PCR was then carried out using a TB Green Premix Ex Taq (Tli RNaseH Plus) kit (Takara, Japan) according to the manufacturer’s protocol. Briefly, 25ng of cDNA, along with 5 μL of 2X TB Green Premix Ex Taq (RNaseH Plus) and 0.2 μL of each 10 μM forward and reverse primer, were added to a total reaction volume of 10 μL. Thermocycling was performed in a CFX96 Touch Real-time PCR Detection System (Bio-Rad Laboratories, Hercules, CA, USA). The primer sequences for *plcR, nprB, nheA, cheY, spo0A, and codY* genes are provided in **Table 1**. Gene expression was normalised to the 16S rRNA housekeeping gene. Data were analysed using the 2^− ΔΔCt^ method and plotted as fold change **(Livak & Schmittgen, 2001)**.

**Table 1.**
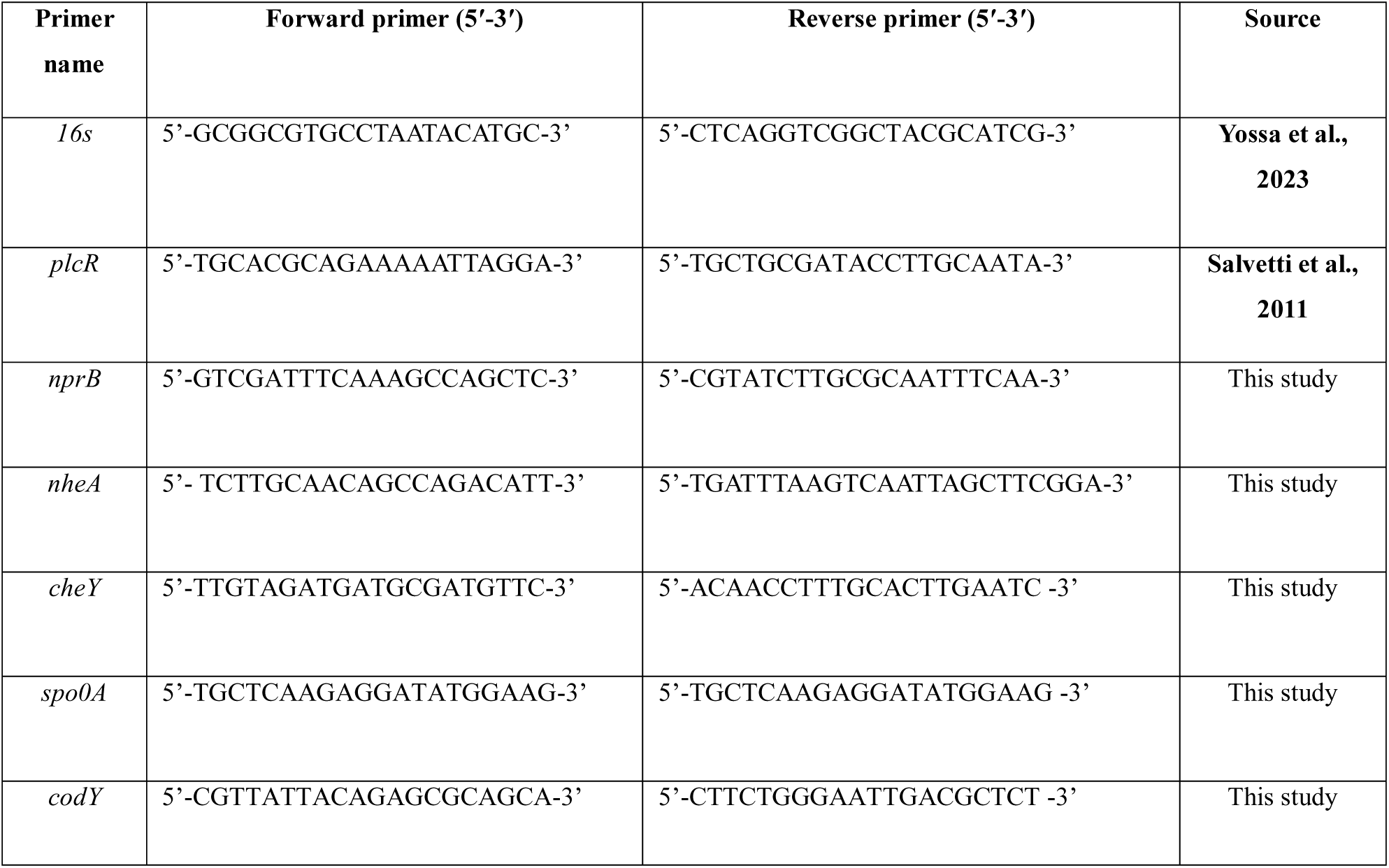
Primers used in the present study.

### 2.9 Statistical analysis

All experiments were performed independently thrice, and the data were expressed as mean±SEM. Microsoft Excel and GraphPad Prism 8.0.1 were used to analyse the data. Statistical significance was analysed using Student’s *t*-test (2-tailed, unpaired). *P-value* less than 0.05 was considered to be statistically significant (\**P-value* ≤ 0.05; \*\**P-value* ≤ 0.01; \*\*\**P*-value ≤ 0.001; ****, *P-value* ≤ 0.0001).

## 3. Results

### 3.1 Antibiotic susceptibility test and determination of Minimum Inhibitory Concentration of gentamicin

The sensitivity of *Bacillus cereus* MSM-S1 to different antibiotics was determined by the octa disc diffusion test. The strain was found to be sensitive to cephalothin, colistin sulphate, gentamicin, streptomycin, and tetracycline (**Figure S1**). Among these antibiotics, gentamicin was selected for our study based on its broad-spectrum activity and common use in treating *Bacillus cereus* infections **(Mills et al., 2022).** Further, the minimum inhibitory concentration (MIC) of gentamicin against *Bacillus cereus* MSM-S1 was calculated and found to be 10 µg/mL (**Figure S2**).

### 3.2 Effect of different doses of gentamicin on the swarming ability of Bacillus cereus MSM-S1

The swarming ability of *Bacillus cereus* MSM-S1 was assayed on nutrient agar plates with 0.6% agar supplemented with 0.25µg/mL (MIC/40) and 0.5µg/mL (MIC/20) of gentamicin. In the absence of gentamicin, the colony exhibited uniform swarming with a self-similar pattern (**Figure S3a**), whereas 0.25µg/mL gentamicin produced a smaller colony with a distinct clear zone around the centre. However, the self-similar pattern was not altered in the presence of 0.25µg/mL gentamicin (**Figure S3b**). In the presence of 0.5µg/mL gentamicin, the colony initially displayed restricted growth and motility; however, from day 5, protrusions started to emerge from the colony and the colony swarmed with disruption of the self-similar pattern (**Figure S3c**). The results are also evident with a reduction in the area covered by the colonies growing in the presence of 0.5µg/mL gentamicin in comparison to the control (**Figure S3d**). Transmission electron microscopy revealed a reduction in flagellation on peripheral cells of *Bacillus cereus* MSM-S1 swarming in the presence of gentamicin in a dose-dependent manner (**Figure S4**). The peripheral cells of *Bacillus cereus* MSM-S1 swarming in the absence and presence of gentamicin were observed through structured illumination microscopy and found to be elongated in the presence of gentamicin (**Figure S5**). Moreover, the mRNA expression of *nheA* was checked in the peripheral cells of the *Bacillus cereus* MSM-S1 colony, swarming in the absence and presence of 0.25µg/mL and 0.5µg/mL gentamicin. It was found that the relative mRNA expression of the *nheA* gene was significantly upregulated to 4.091 ± 0.916-fold in the presence of 0.25µg/mL gentamicin and to 2.51 ± 0.441-fold in the presence of 0.5µg/mL gentamicin, as compared to the control (**Figure S6**). Since gentamicin at a concentration of 0.5μg/mL showed a greater effect on colony morphology as well as colony movement, we studied the impact of the presence of a sibling colony of *Bacillus cereus* MSM-S1 in the presence and absence of 0.5μg/mL concentration of gentamicin.

### 3.3 Sibling colonies of Bacillus cereus MSM-S1 showed rivalry when spotted adjacent to each other

The sibling colonies of *Bacillus cereus* MSM-S1 were allowed to interact by spotting 2µL of *Bacillus cereus* MSM-S1 culture (10^7^ CFU/mL) adjacent to each other on semi-solid nutrient agar plates with 0.6% agar in the absence and presence of a subinhibitory concentration (0.5µg/mL) of gentamicin. When *Bacillus cereus* MSM-S1 was spotted alone on the plates without gentamicin, swarming of the colony was observed over days with a symmetrical colony morphology representing a self-similar pattern (**Figure 1a**). During the interaction between sibling colonies in the absence of gentamicin, the two colonies initially swarmed with a symmetrical colony morphology and growth. However, from day 5 onwards, the symmetry of the interacting colonies was observed to be altered at the edge where the two colonies faced each other (interacting zone). On day 7, the symmetry of the interacting colonies at the interacting zone was completely altered, forming a clear line of demarcation or a zone of no growth between the sibling colonies (**Figure 1b**).

**Figure 1.**
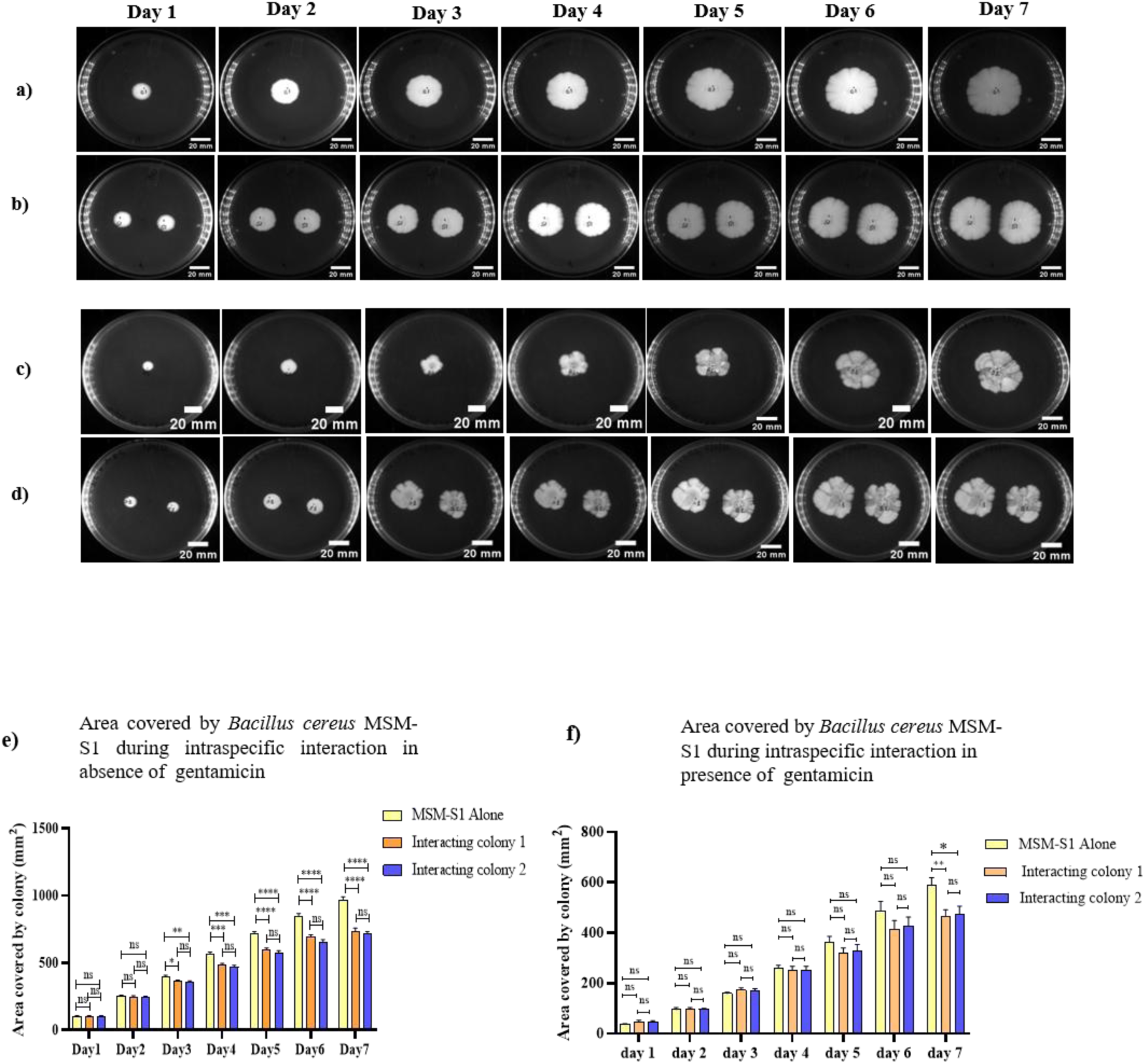
Representative images of (a) *Bacillus cereus* MSM-S1 growing alone in the absence of gentamicin (b) Sibling colonies of *Bacillus cereus* MSM-S1 growing adjacent to each other at 3cm in absence of gentamicin (c) *Bacillus cereus* MSM-S1 growing alone in the presence of gentamicin (d) Sibling colonies of *Bacillus cereus* MSM-S1 growing adjacent to each other at 3cm in the presence of gentamicin (e) Area covered by colonies during interaction in the absence of gentamicin (f) Area covered by colonies during interaction in the presence of gentamicin. Data presented as the mean ± SEM (\**p-value* ≤ 0.05; \*\**p-value* ≤ 0.01; \*\*\**p* value ≤ 0.001; ****, *p-value* ≤ 0.0001).

While growing alone in the presence of gentamicin, the swarming pattern of the colony was altered, with protrusions emerging from the colony and an altered self-similar pattern (**Figure 1c**). During interaction in the presence of gentamicin, the alteration in the symmetry of the colonies at the interacting zone was observed and by day 7, the demarcation line formed between the colonies (**Figure 1d**). Furthermore, reduction in the area covered by the interacting colonies with respect to control (colony growing alone) colonies, both in the absence and presence of gentamicin (**Figure 1e and 1f**) was observed.

Moreover, the radius from the center of the colony to the non-interacting and interacting edge of the colony was measured, and it was observed that both in the absence and presence of gentamicin, the radius towards the interacting zone starts to decrease significantly from day 4 onwards (**Figure S7**), suggesting decreased motility of the colony towards the interacting zone.

Based on the results of colony movement and morphology, a time-lapse study was conducted on three different day points to further understand the cellular-level dynamics, such as the arrangement and surface topology of peripheral cells from the colony growing alone, the non-interacting zone and the interacting zone.

For further studies, the day points were selected based on the colony morphology as follows:

- **Day 3**: the day when no change in the symmetry of the interacting colonies was observed.
- **Day 5**: the day when the symmetry of the interacting colonies was observed to be altered.
- **Day 7**: the day after the symmetry of the interacting colonies was altered and a clear line of demarcation formed between the colonies.

### 3.4 Intraspecific interaction between sibling colonies of Bacillus cereus MSM-S1 leads to alteration in the arrangement of peripheral cells at the interacting zone (IZ)

Structured illumination microscopy was performed to gain insights into the day-wise arrangement of peripheral cells of the *Bacillus cereus* MSM-S1 during intraspecific interaction between sibling colonies in the absence and presence of gentamicin. In the absence of gentamicin, the peripheral cells of the colony growing alone and in the non-interacting zone were observed to be arranged in long, elongated chains in a swirled pattern on days 3, 5 and 7 (**Figure 2a)**. However, in the interacting zone, the cells showed a gradual alteration in their arrangement over the course of the study. On day 3, the cells were arranged in a swirled fashion, as observed in the colony swarming alone. On day 5, this arrangement was found to be disrupted, and the cells were no longer arranged in the swirling pattern. Furthermore, on day 7, the swirl pattern of arrangement was completely lost, with cells randomly arranged, indicating cell jamming (**Figure 2a**). In the presence of gentamicin, the cells in the colony growing alone and in non-interacting zone were arranged as chains (**Figure 2b**); however, the swirling pattern appeared different from that of the colony growing in the absence of gentamicin. At the interacting zone, the cells were initially arranged as chains (day 3) but began to lose this arrangement after day 5, and the swirl like pattern was lost by day 7 (**Figure 2b**). It is noteworthy that on day 7, the cells were not as randomly arranged as observed in the absence of gentamicin, and the cell jamming was less.

**Figure 2.**
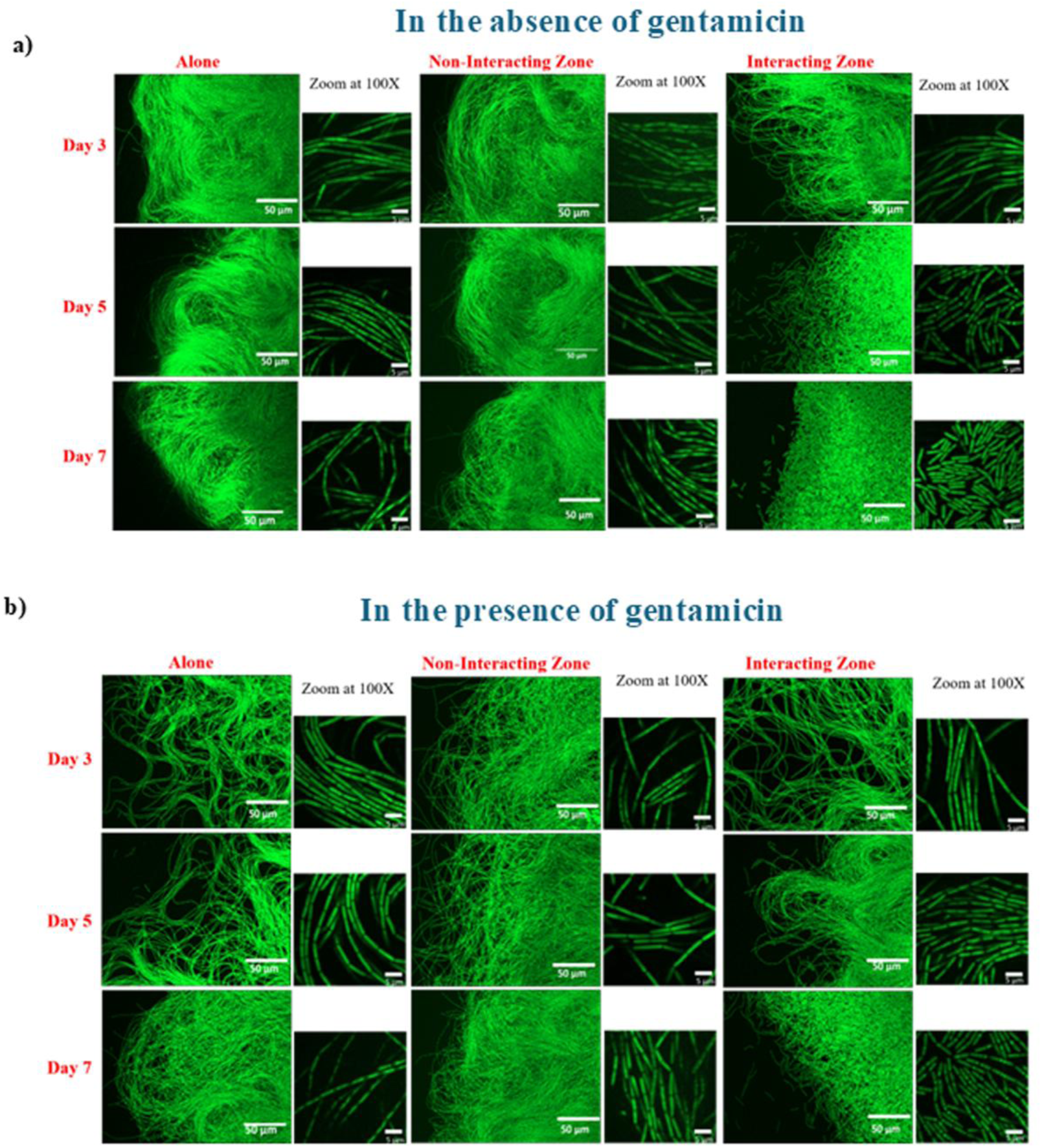
Arrangement of cells from the colonies growing alone, non-interacting, and the interacting zone a) in the absence and b) in the presence of gentamicin. Images taken at 40X magnification; zoom images taken at 100X magnification. Images represented from three independent biological replicates, N=3. The white bar represents 50µm in 40X images and 5µm in 100X magnified images.

### 3.5 Intraspecific interaction between sibling colonies of Bacillus cereus MSM-S1 leads to alteration in the ultrastructure of peripheral cells at the interacting zone (IZ)

To further examine the surface morphology and cell architecture of *Bacillus cereus* MSM-S1 during intraspecific interactions between sibling colonies, in the absence and presence of gentamicin, field-emission scanning electron microscopy was performed. In the presence and absence of gentamicin, the peripheral cells of the colony growing alone and the non-interacting zone appeared rod-shaped and showed little change over the days of the study. However, striking differences were observed in the cells at the interacting zone. In the absence of gentamicin, on day 3 and day 5, the morphology of the cells at the interacting zone had altered, and a few cells showed early sporulation, which on day 7 was observed to have completely formed spores (**Figure 3a**). In the presence of gentamicin, the cells showed elongation while growing alone, both in the non-interacting and in the interacting zones. However, observable sporulation was not seen in the cells from colonies growing alone, as well as in the non-interacting zone in the presence of gentamicin. Interestingly, cells at the interacting zone showed the presence of some sporulating cells on day 7 (**Figure 3b**), but the number of sporulating cells was much less as compared to that observed in the interacting zone on day 7 in the absence of gentamicin (**Figure 3a**).

**Figure 3.**
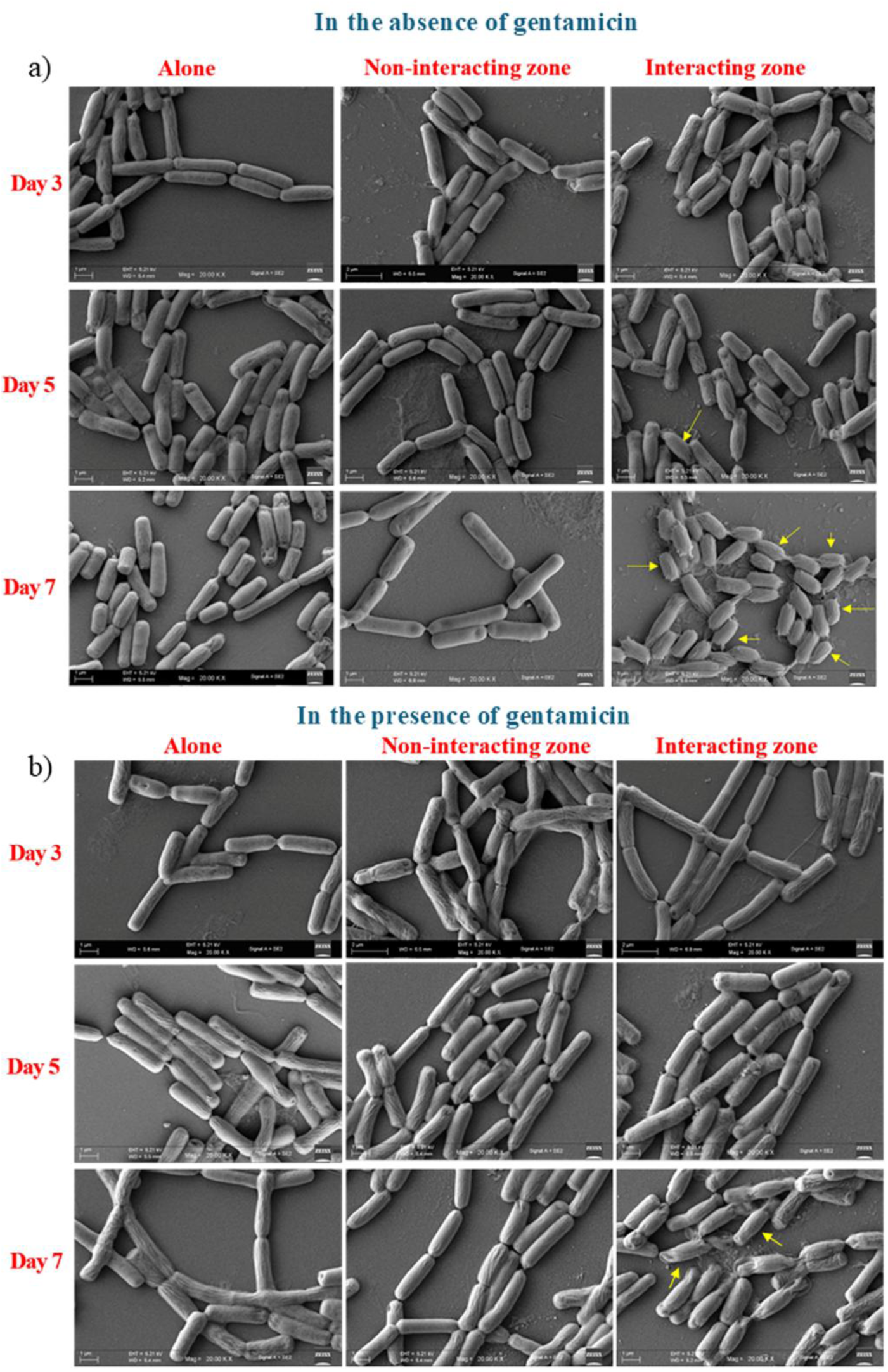
Representative images taken at 20KX magnification showing the ultrastructure of cells from the periphery of colonies growing alone, the non-interacting zone and the interacting zone in the (a) absence of gentamicin and (b) presence of gentamicin. Arrow indicates sporulation of cells.

### 3.6 Nutrient depletion was the major contributory factor behind the formation of the interacting zone

The zone of inhibition between the sibling colonies could have resulted from inhibitory molecule(s) secreted by interacting colonies, which could act specifically on sibling cells, or due to nutrient depletion in the interacting zone. Hence, further experiments were done to find the possible cause behind the formation of the zone of inhibition during intraspecific interaction between sibling colonies

At first, it was assessed whether the interaction zone supports further growth of bacteria of the same and different species when spotted on it at different days of interaction. For this, 2 µL of *Bacillus cereus* MSM-S1 (10^7^ CFU/mL), *Bacillus thuringiensis* KPWP1 (10^7^ CFU/mL) and *Pseudomonas aeruginosa* KPW1.S1 (10^7^ CFU/mL) were spotted between the interacting sibling colonies of *Bacillus cereus* MSM-S1 on day 3, day 5, and day 7 of interaction.

When *Bacillus cereus* MSM-S1 was spotted on the interacting zone on day 3 of interaction, faint colony growth of the newly added *Bacillus cereus* MSM-S1 was observed in the interacting zone. When spotted on day 5 of interaction, the growth of the newly added *Bacillus cereus* MSM-S1 was much less. Further, when spotted on day 7 of interaction, the growth of newly added *Bacillus cereus* MSM-S1 was completely absent in the interacting zone (**Figure 4a**). When *Bacillus thuringiensis* KPWP1 was spotted on the interacting zone on day 3 of interaction, faint colony growth of the newly added *Bacillus thuringiensis* KPWP1 was observed in the interacting zone. When spotted on day 5 and day 7 of interaction, the growth of the newly added *Bacillus thuringiensis* KPWP1 was completely absent in the interacting zone (**Figure S8a**). However, when *Pseudomonas aeruginosa* KPW1.S1 was spotted on interacting zone on day 3, colony growth was observed in the interacting zone. When spotted on day 5 and day 7 of interaction, the growth of *Pseudomonas aeruginosa* KPW1.S1 colony was observed, but the growth was substantially less (**Figure S8b**).

**Figure 4.**
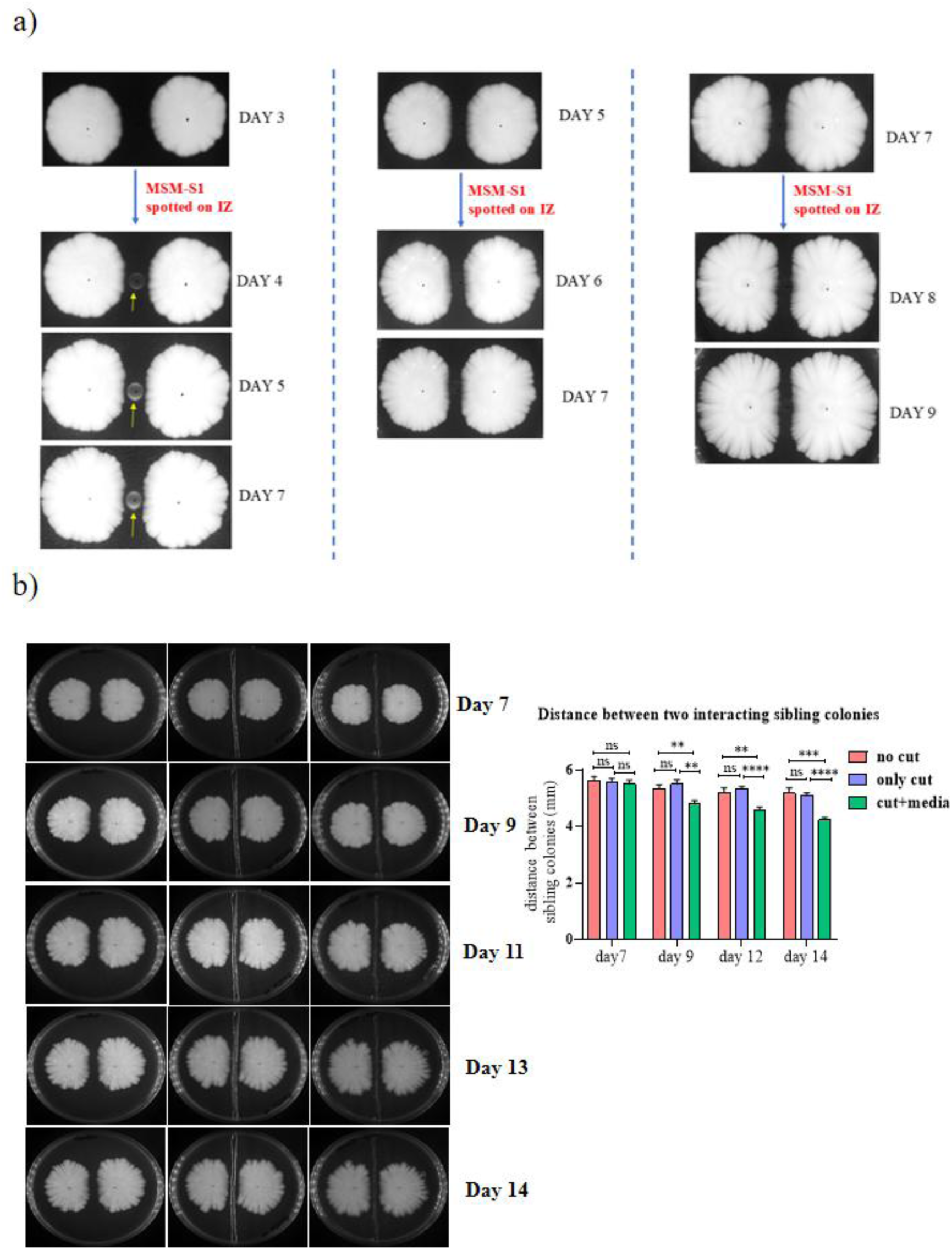
(a) Representative images showing the growth of colonies of *Bacillus cereus* MSM-S1 spotted on the interacting zone on day 3, day 5, and day 7. (b) Growth and movement of the colony at the interacting zone. Data presented as the mean ± SEM, where * is *P* ≤ 0.05, ** is *P* ≤ 0.01, *** is *P* ≤ 0.001, **** is *P* ≤ 0.0001, and ns signifies non-significant.

To further explore the cause behind the formation of the zone of no growth, in another experiment, *Bacillus cereus* MSM-S1 was grown adjacent to each other on nutrient agar plates with 0.6% agar. Upon observing the pattern breaks in the interacting colonies on day 7, the agar in the interacting zone was cut to make a narrow rectangular well and filled with fresh media and the interacting colonies were allowed to grow further. It was observed that the distance between two interacting sibling colonies was reduced in plates that were filled with fresh media (**Figure 4b**), which indicates restoration of colony movement in the presence of fresh growth medium. This suggested the depletion of nutrients at the interacting zone during the interaction between sibling colonies of MSM-S1, which was compensated by the addition of fresh media at the interacting zone; thus, it signifies the possibility of contribution of nutrition limitations in regulating the zone of inhibition between two interacting sibling colonies of MSM-S1.

To further confirm whether nutrient limitation was really playing the major role behind formation of the demarcation line, sibling colonies were grown adjacent to each other on nutrient agar plates with 0.6% agar with increased peptone concentrations at varying distances (1cm, 2cm, 3cm). It was observed that the sibling colonies could merge when spotted at a distance of 1 cm on plates with increased peptone concentrations, which further indicated that the intraspecific interaction between the colonies was being modulated by nutrient levels at the interacting zone (**Figure S9a**). As the previous results indicated that the nutrient level governs the formation of a zone of inhibition between two sibling colonies of MSM-S1 and two growing colonies merge when spotted 1 cm apart from each other, microscopic analyses of the arrangement of the cells at the non-interacting and interacting zones during intraspecific interaction between sibling colonies growing at 1cm distance on semi-solid nutrient agar plates with increased peptone concentrations were performed. It was observed that the cells at the non-interacting zone from the control (normal peptone concentration), 2X peptone, 6X peptone, and 8X peptone displayed a swirling arrangement. Interestingly, no distinct edge of cells was observed in the interacting zone, indicating merging of two colonies at the interacting zone, and increased cell densities were observed with increased peptone concentrations for both the interacting and non-interacting zones (**Figure S9b**).

### 3.7 Differential expression of genes in Bacillus cereus MSM-S1 during intraspecific interaction in the absence and presence of gentamicin

To examine if the observed differences in the peripheral cells at the interacting zone at the cellular level was mediated through differential gene expressions, the expression of genes related to quorum sensing (*plcR, nprB*), virulence (*nheA*), sporulation (*spo0A*), chemotaxis (*cheY*), and, a pleiotropic transcriptional regulator that affects the expression of genes based on nutritional availability, *codY,* was studied in the peripheral cells of the *Bacillus cereus* MSM-S1 colony growing alone, in non-interacting zone, and in the interacting zone during intraspecific interaction in the absence and presence of gentamicin. It was observed that in the absence of gentamicin, as compared to the control (i.e. the colonies growing alone), the relative mRNA expression of the *plcR* gene was reduced to 0.307±0.165-fold in the cells at the interacting zone, while no significant change was observed in the cells at the non-interacting zone (**Figure 5a**), suggesting a downregulation of quorum sensing at the interacting zone. In the presence of gentamicin, the relative mRNA expression of *plcR* upregulated to 3.034±0.347-fold and *4*.58±1.10-fold at the non-interacting and interacting zones, respectively (**Figure 5a**), as compared to colonies growing alone. The expression of the *nprB* gene was observed to be downregulated, with relative expression reduced to 0.67 ± 0.068-fold in the non-interacting zone and further decreased to 0.447 ± 0.099-fold in the interacting zone compared to colonies growing alone in the absence of gentamicin (**Figure 5b).** In the presence of gentamicin, an increase in expression to 1.421±0.176 fold in the non-interacting zone was observed, which was further increased in the interacting zone (1.936±0.174 fold) compared to colonies growing alone in the presence of gentamicin (**Figure 5b).**

**Figure 5.**
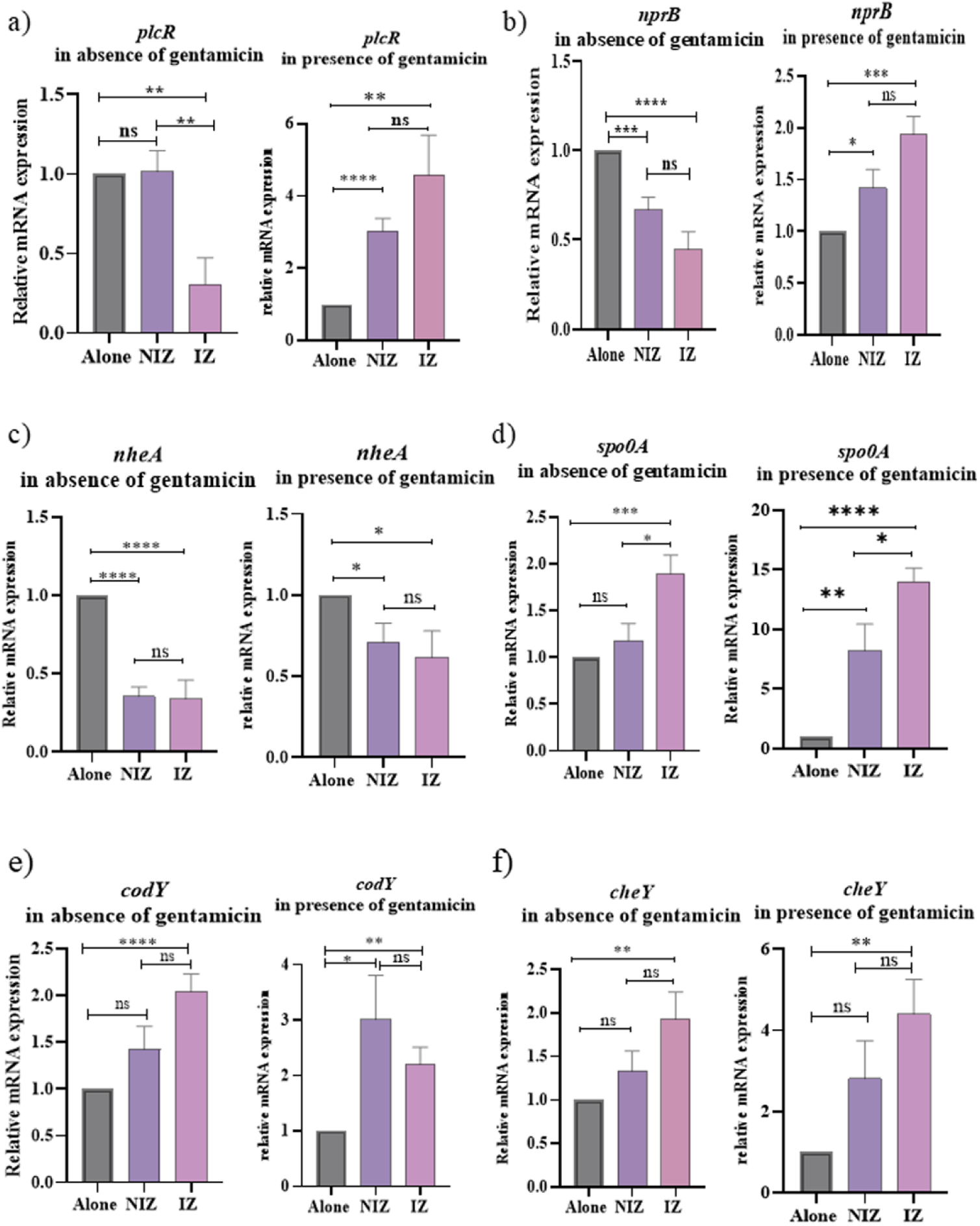
Differential expression of (a) *plcR,* (b) *nprB,* (c) *nheA,* (d) *spo0*A, (e) *codY, and* (f) *cheY* genes in *Bacillus cereus* MSM-S1 during intraspecific interaction in the absence and presence of gentamicin. Data presented as the mean ± SEM, where * is *P* ≤ 0.05, ** is *P* ≤ 0.01, *** is *P* ≤ 0.001, **** is *P* ≤ 0.0001, and ns signifies non-significant.

The expression of *nheA*, which codes for the “A” component of the tripartite non-hemolytic enterotoxin, was also reduced in the cells at the interacting zone in both the absence and presence of gentamicin. In the absence of gentamicin, the expression was reduced to 0.340±0.119-fold, while in the presence of gentamicin, it was reduced to 0.617±0.162-fold compared to colonies growing alone in the absence and presence of gentamicin, respectively (**Figure 5c).**

The cells at the interacting zone were observed to have increased expression of the *spo0A* gene in both the absence and presence of gentamicin. In the absence of gentamicin, the expression at the interacting zone was upregulated to 1.894±0.199-fold; in the presence of gentamicin, it was upregulated to 14.012±1.11-fold compared to colonies growing alone in the absence and presence of gentamicin, respectively. In the presence of gentamicin, an upregulation to 8.213±2.224-fold was also observed in cells at the non-interacting zone (**Figure 5d**).

Strikingly, the relative mRNA expression of *codY* was increased in the cells at the interacting zone both in the presence and absence of gentamicin. In the absence of gentamicin, a 2.046±0.187-fold increase, while in the presence of gentamicin, a 2.206±0.304-fold increase was observed in cells at the interacting zone, indicating a possible interplay of molecular pathways functioning based on the nutritional status of the cells (**Figure 5e**). However, an increase in the expression of codY to 3.011±0.798-fold was also observed in cells at the non-interacting zone in the presence of gentamicin. Interestingly, the expression of *cheY* in peripheral cells of the interacting zone was observed to be upregulated to 1.99±0.405-fold and 4.397±0.855-fold compared to colonies growing alone in the absence and presence of gentamicin, respectively (**Figure 5f**).

## 4. Discussion

The present study investigates the impact of intraspecific interaction between sibling colonies of *Bacillus cereus* MSM-S1 growing on semi-solid nutrient agar plates (0.6% agar) in the absence and presence of gentamicin. *Bacillus cereus* MSM-S1, when spotted alone in the absence of gentamicin in the growth medium, exhibited swarming with uniform and symmetric colony morphology, but in the presence of gentamicin, an initial restriction in colony growth was observed but later protrusions of colony growth appeared and the cell were found to be elongated with less flagellation and increased expression of toxin gene *nheA*. However, interaction between the sibling colonies markedly altered their morphology and reduced their movement towards the interacting zone, which manifested as the formation of a distinct line of demarcation between them. Such instances of sibling colony interaction have been reported previously, where colonies are prevented from intruding on each other’s niche. Cruz-López R et al. showed sibling rivalry in *Marinobacter algicola* caused by loss of motility of cells at the interacting face **(Cruz-López et al., 2021)**. Similar rivalry among sibling colonies of bacteria has also been observed in swimming colonies of *Bradyrhizobium diazoefficiens* **(Sandhu et al., 2025)**, which occurred due to suppression of bacterial growth during the interaction.

When the sibling colonies were made to interact in the presence of gentamicin, the swarming of *B. cereus* MSM-S1 was reduced, with altered colony morphology, which suggested that gentamicin acted as a stress factor for the swarming colonies. Interestingly, upon interaction with the sibling colonies, the colonies showed a similar effect as in the absence of gentamicin, with a distinct line of demarcation forming between the interacting colonies. Surprisingly, even in the presence of antibiotics in the growth medium, which is a common threat to both colonies, the interacting colonies did not co-mingle to increase their population to cope with the antibiotics; rather, they remained separated and displayed restricted motility towards each other.

Microscopy results revealed that, while on the initial days of interaction (day 3), the peripheral cells at both the non-interacting and interacting edges displayed uniformity in their pattern (as associated chains), as observed in the control (colonies growing alone), this uniformity was lost in cells at the interacting zone over time. In the absence of gentamicin, on day 7, the cells at the interacting zone appeared to be randomly arranged and had lost their association as chains. This indicated that, as the sibling colonies were growing and the intercolony distance reduced over days, unfavourable conditions were being imposed on the cells, and as a result, the cells lost their homogeneity in arrangement as chains and displayed cell jamming. A similar pattern of change was observed when colonies were growing in the presence of gentamicin; however, the cells exhibited less jamming on day 7, suggesting that in the presence of gentamicin, the cells preferred to stay associated to overcome the dual stress of antibiotic and a sibling colony

Scanning electron microscopy images depicted a striking difference in the morphology of the peripheral cells from the colony growing alone, non-interacting, and interacting zones. The cells at the interacting zone appeared to be highly stressed on day 7, and the cells had sporulated. On the other hand, the number of spore cells at the periphery of the colony growing alone and in the non-interacting zones was much less on day 7. Sporulation is one of the most advantageous adaptations acquired by *Bacillus sp*. **(De Vries et al., 2004)** and ensures the persistence of *Bacillus* in harsh environments. This indicated that the presence of a sibling colony of *Bacillus cereus* MSM-S1 in the vicinity was contributing to the stress experienced by the cells at the interacting zone and that *Bacillus cereus* MSM-S1 cells employed sporulation as a strategy to survive under stress imposed on them due to the presence of the sibling colony. However, it was noteworthy that in the presence of gentamicin, the cells did not form spores while growing alone and in the non-interacting zone. Interestingly, sporulation was observed only in the interacting zone, but the extent of sporulation was less in the presence of gentamicin as compared to that in the absence of gentamicin. This could have been because gentamicin is an inhibitor of protein synthesis in bacteria; the overall process of sporulation was impacted in the cells, and hence less sporulation cells were observed.

The microscopic results revealed alterations in the cellular arrangement and morphology of the cells at the interacting zone. The formation of spores further indicated that the cells at the interacting zone experienced stress during interaction with a sibling colony. All these observations raised questions regarding the cause behind the formation of the zone of inhibition between the interacting sibling colonies of *Bacillus cereus* MSM-S1. The zone of inhibition between the sibling colonies could have resulted from inhibitory molecule(s) secreted by interacting colonies, which could act specifically on sibling cells, or due to nutrient depletion in the interacting zone. Hence, further experiments were done to find the possible cause behind the formation of the zone of inhibition during intraspecific interaction between sibling colonies. Our experiments suggested that once the line of demarcation forms (day 7), the interacting zone could not support the growth of fresh inoculum of *Bacillus cereus* MSM-S1 or *Bacillus thuringiensis* KPWP1 spotted on it. This indicated that there could either be depletion of nutrients or diffusible inhibitor molecule(s) that might have accumulated in the interacting zone. However, *Pseudomonas aeruginosa* KPW1.S1 spotted on the interaction zone could still grow but the growth was less compared to when spotted on day 3. This could have happened either due to production or accumulation of inhibitor(s) or compounds released by *Bacillus cereus* MSM-S1 that restricted the growth of *Pseudomonas* or due to nutrient limitation at the interacting zone. Lower growth of *Pseudomonas* when spotted on day 7, but no growth of *Bacillus cereus* MSM-S1 or *Bacillus thuringiensis* KPWP1, also indicates different nutritional requirements of two different genera of bacteria.

The observation of the reduction in the distance between the two sibling colonies upon addition of fresh media (nutrients) at the interacting zone further indicated nutrient limitation between the interacting sibling colonies of *Bacillus cereus* MSM-S1. It was also observed that the interacting colonies could merge when grown at 1cm distance on nutrient agar plates with increased peptone concentrations. The results indicated that nutrients play a major role in the formation of the line of demarcation at the interacting zone, although we cannot completely deny or rule out the involvement of inhibitory molecule(s) in the formation of the zone of no growth between the interacting sibling colonies.

Since the alterations in the cellular behaviour of bacteria are a reflection of altered expression of genes that help the bacteria to adapt to their external environment, the differential expression of several genes was studied to comprehend their roles in the cellular response during the intraspecific interaction between the sibling colonies of *Bacillus cereus* MSM-S1.

Signals, such as molecular cues and stimuli, are essential for bacterial communication and information exchange, allowing bacteria to sense and react to their environment **(Liu et al., 2025)**. The cross-talk between the same or different species of bacteria is mediated through quorum sensing, which involves the release and sensing of small diffusible signaling molecules **(Diggle et al., 2007; Keller & Surette, 2006).** The expression of the *plcR* gene encoding the quorum sensing master regulator PlcR in *Bacillus cereus* regulates several genes encoding proteins associated with cell defence, environmental sensing, and virulence (phospholipases, proteases, haemolysins, toxins, etc.), thereby helping cell survival **(Gohar et al., 2008; Qin et al., 2025)**. Since the behavioural and physiological changes in bacteria greatly rely on quorum sensing **(Rutherford & Bassler, 2012)**, the expression of quorum sensing related genes (*plcR, nprB*) was studied. In the absence of gentamicin, the cells at the interacting zone exhibited a significant decrease in the expression of quorum-sensing genes, which accounted for the loss of cellular arrangement observed on day 7 of interaction, as observed in microscopic analysis (**Figure. 2a**). Strikingly, the expression of quorum sensing genes was observed to be upregulated in the presence of gentamicin (**Figure. 5a and 5b**). This could have happened because gentamicin itself is a stress to the growing cells; to grow and swarm, it modulated its quorum-sensing mechanism to remain associated with each other to survive in such conditions.

It was observed that the expression of the spo0A gene, which encodes the regulator of sporulation Spo0A **(Hoch, 1993)** increased in cells at the interacting zone, both in the absence and presence of gentamicin, which supported the microscopic observation of sporulation in this zone (**Figure. 5d**). Interestingly, the expression of the virulence gene *nheA* was observed to be reduced in the interacting zone both in the absence and presence of antibiotic, which indicates that availability of resources regulates the decision-making in MSM-S1 cells to make or not to make the toxin as a trade-off between making the toxin and forming spores.

The interesting phenomenon observed in our study was the upregulation of the c*odY* gene in the cells at the interacting zone (**Figure. 5e**). CodY is a pleiotropic global transcriptional regulator that regulates gene expression in response to environmental nutrient limitation. CodY functions as an active transcriptional repressor that binds to DNA during rapid exponential growth and is liberated from DNA during the shift into the stationary phase, allowing the expression of genes involved in nutrient limitation adaptation **(Geiger & Wolz, 2014)**. It has been shown that the transcription of numerous genes involved in *B. subtilis* competence, sporulation, and biofilm formation is inhibited by CodY **(Lindbäck et al., 2012)**. In their report, Lindbäck et al. have also demonstrated the downregulation of expression of flagellar and motility genes in the Δ*codY* mutant, which suggests the role of CodY in regulating motility as an adaptation of cells during starvation. Although CodY has been reported to inhibit sporulation, some reports suggest its role as a positive regulator of *spo0A* expression. Kim et al. reported a downregulation of sporulation regulator genes by the CodY mutation during glucose starvation **(Kim et al., 2016)**. Another report by Mirouze et al also suggests a similar positive effect of CodY on the expression of *spo0A* **(Mirouze et al., 2011)**. These studies corroborate the observed upregulation in the expression of both *codY* and *spo0A* in the cells at the interaction zone, indicating the role of *codY* in regulating sporulation under nutrient starvation during interaction between sibling colonies of *Bacillus cereus* MSM-S1.

Another interesting observation was the upregulation of the chemotaxis response regulator *cheY* gene in the cells at the interacting zone (**Figure. 5f**). Motile bacteria can move towards more favourable conditions by sensing their chemical environment through chemotaxis **(Tang et al., 2010)** and regulating the direction of flagellar rotation. An increased expression of the *cheY* gene in the cells at the interacting zone indicates that cells are experiencing a strong chemotactic signal as a stress response to sense and respond to the local environment. Thus, the study paves the way for future research on the roles of *codY* and *cheY* to decipher their roles during intraspecific interaction between sibling colonies of *Bacillus cereus*.

## 5. Conclusion

To conclude, the present study reveals that although the presence of gentamicin could reduce swarming of *Bacillus cereus* MSM-S1 and altered its colony morphology, the presence of a sibling colony appeared to be a greater challenge, as the motility was restricted at the interacting zone where two sibling colonies faced each other. The cells of *Bacillus cereus* MSM-S1 tried to stay connected and reduced cell jamming at the interaction zone in the presence of gentamicin, indicating a survival strategy to thrive under the stress of gentamicin as well as presence of a sibling colony. It was also revealed that nutrient limitation in the interacting zone is a major contributory factor towards the formation of a zone of no growth between the interacting sibling colonies of *Bacillus cereus* MSM-S1. The study reveals a strategic adaptation employed by the cells to survive in a competitive environment through the modulation of cell length, toxin production, sporulation and quorum sensing as survival strategies employed by *Bacillus cereus* MSM-S1 to cope with the stress conditions. The present study also opens future avenues to explore and elucidate the role of chemotaxis, the role of the *codY* gene and other contributing factors that help *Bacillus cereus* MSM-S1 to perceive the presence of a sibling colony in the absence and presence of gentamicin, to understand different survival mechanisms employed by *Bacillus cereus* under stress conditions that would help in better therapeutic interventions.

## Acknowledgements

KP acknowledges the Department of Biotechnology (DBT), India, for providing a research fellowship. The authors thank Mr Ritabrata Ghosh for structured illumination microscopy and Mr Kasinath Sahu for Scanning Electron Microscopy

## Author Contributions

KP conceptualized, designed, executed the study, analyzed the data and written the manuscript. BC isolated and characterized the bacterial strain, SKS analyzed the data and edited the manuscript. TKS conceptualized, designed and supervised the study, analyzed the data and edited the manuscript.

## Availability of Data and Materials Declaration

All data generated or analyzed in this study are included in the article.

## Ethical Approval

Not applicable.

## Consent to Participate

All authors had consented to participate in the study.

## Consent for Publication

All authors have given consent for publication.

