## Supplementary Data for "Exploring the impact of intraspecific interaction between sibling colonies of *Bacillus cereus* MSM-S1 in the absence and presence of antibiotics"

^*^ORCID ID- 0000-0001-6482-914X

**Figure S1**

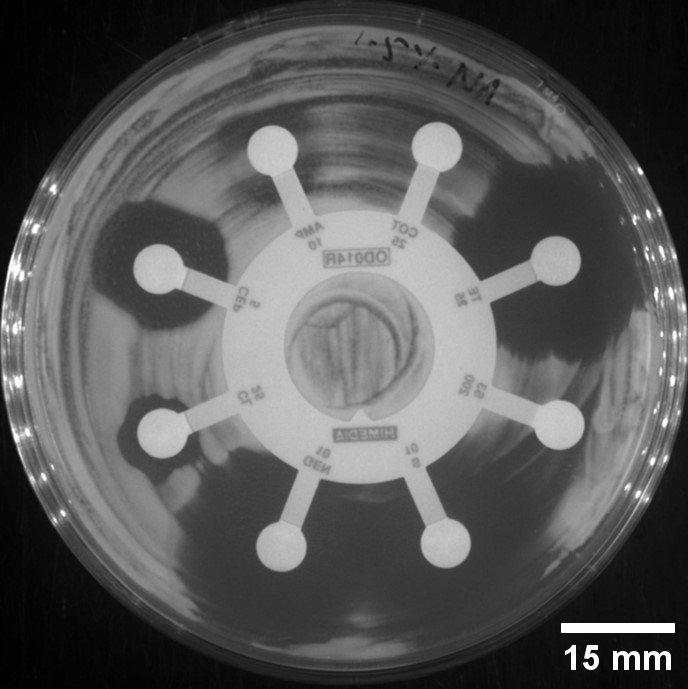

**Figure S1.** Sensitivity of *Bacillus cereus* MSM-S1 to different antibiotics. The circle represents gentamicin. Antibiotic disc used: HiMedia OD014R-1PK G-IV-minus.

**Figure S2**

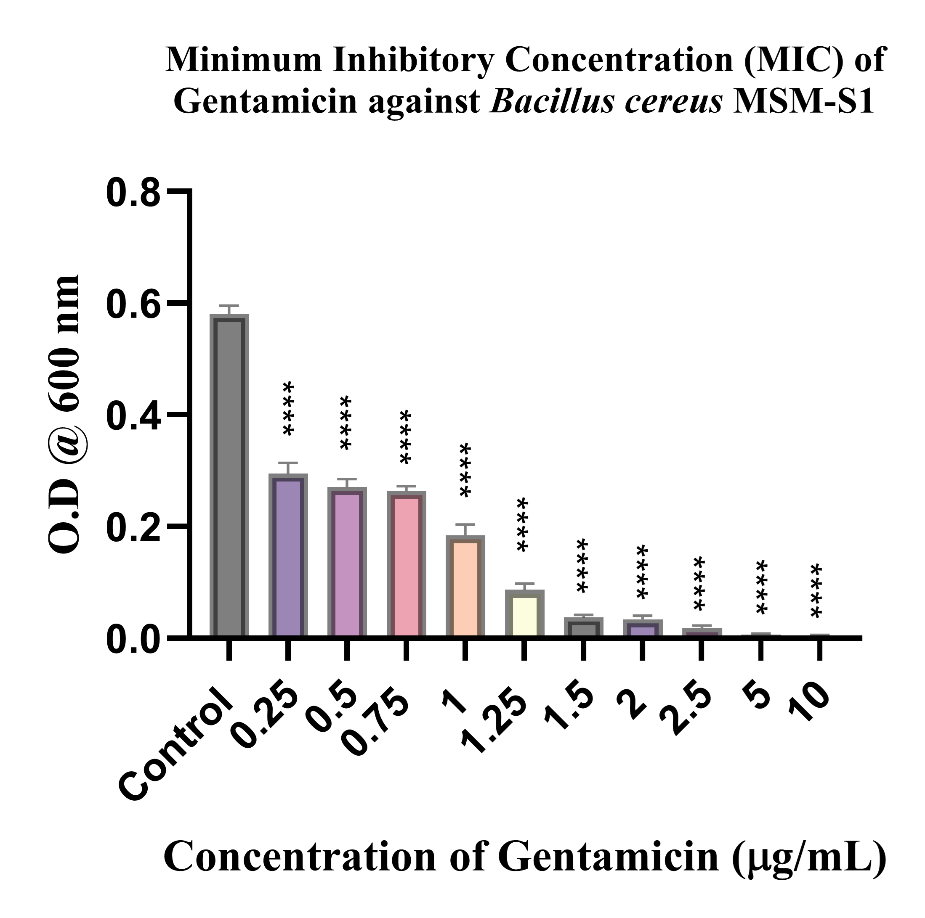

**Figure S2.** Minimum Inhibitory Concentration of Gentamicin against *Bacillus cereus* MSM-S1. Data presented as the mean ± SEM where * is *P* ≤ 0.05, ** is *P* ≤ 0.01, *** is *P* ≤ 0.001, **** is *P* ≤ 0.0001, and ns signifies non-significant.

**Figure S3**

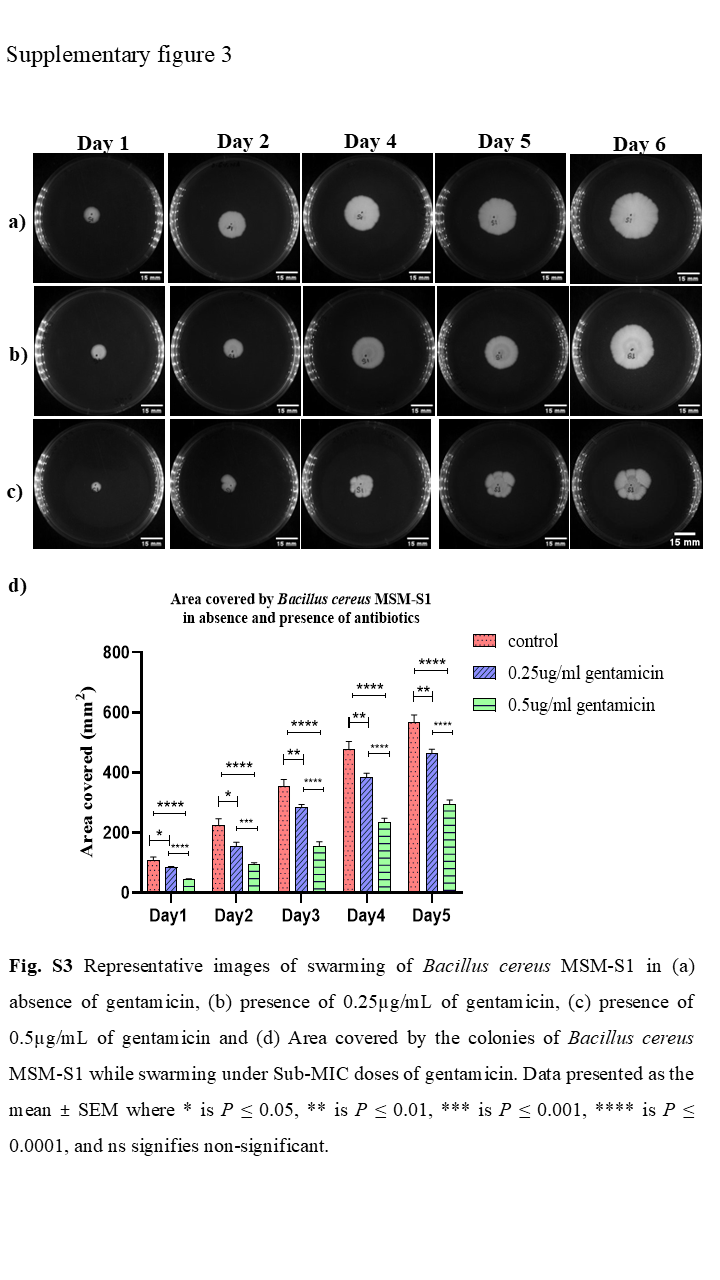

**Figure S3.** Representative images of swarming of *Bacillus cereus* MSM-S1 in (a) absence of gentamicin, (b) presence of 0.25µg/mL of gentamicin, (c) presence of 0.5µg/mL of gentamicin and (d) Area covered by the colonies of *Bacillus cereus* MSM-S1 while swarming under Sub-MIC doses of gentamicin. Data presented as the mean ± SEM where * is *P* ≤ 0.05, ** is *P* ≤ 0.01, *** is *P* ≤ 0.001, **** is *P* ≤ 0.0001, and ns signifies non-significant.

**Figure S4**

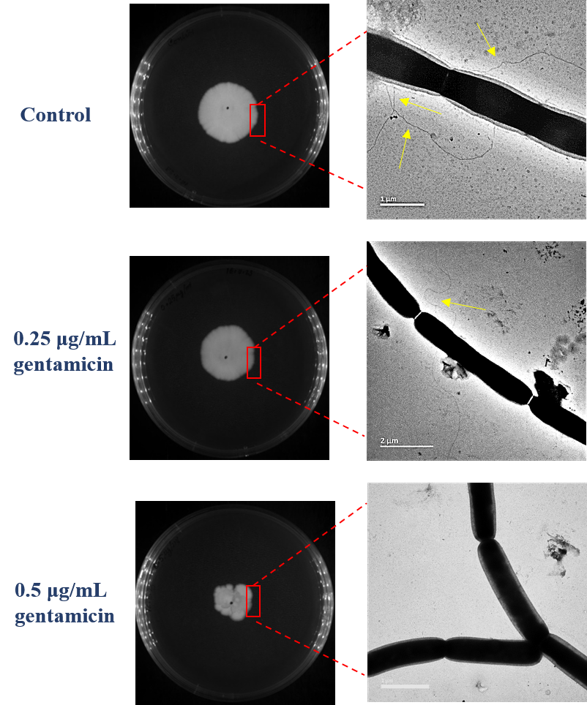

**Figure S4.** Flagellation of *Bacillus cereus* MSM-S1 cells during swarming in the absence and presence of gentamicin on Day 5, N=3. The yellow arrow indicates flagella on the cells.

**Figure S5**

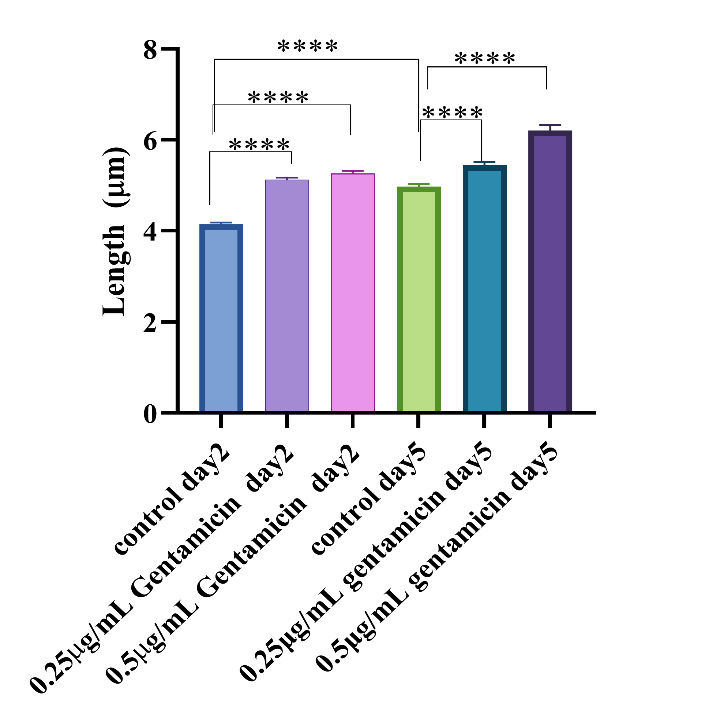

**Figure S5.** Length of cells during swarming under antibiotics on two different days. Data presented as the mean ± SEM for N=3, where * is *P* ≤ 0.05, ** is *P* ≤ 0.01, *** is *P* ≤ 0.001, **** is *P* ≤ 0.0001, and ns signifies non-significant.

**Figure S6**

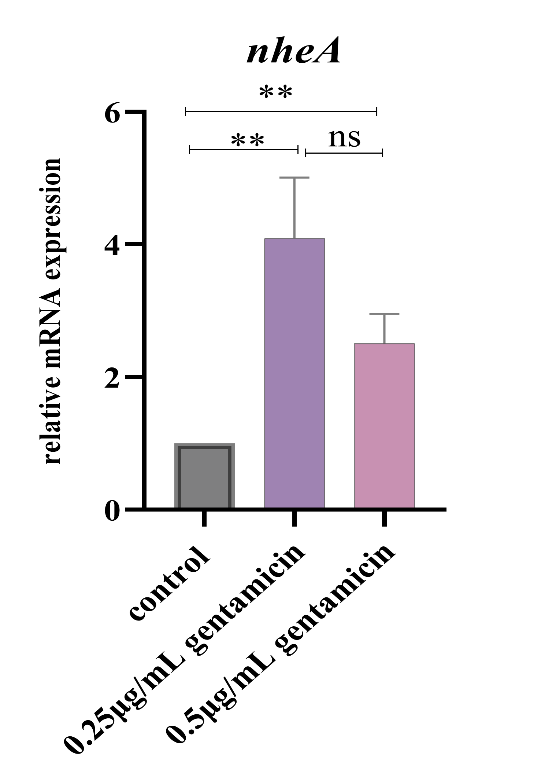

**Figure S6.** Expression of *nheA* in *Bacillus cereus* MSM-S1 during swarming in the absence and presence of gentamicin. Data presented as the mean ± SEM for three independent biological replicates, where * is *P* ≤ 0.05, ** is *P* ≤ 0.01, *** is *P* ≤ 0.001, **** is *P* ≤ 0.0001, and ns signifies non-significant.

**Figure S7**

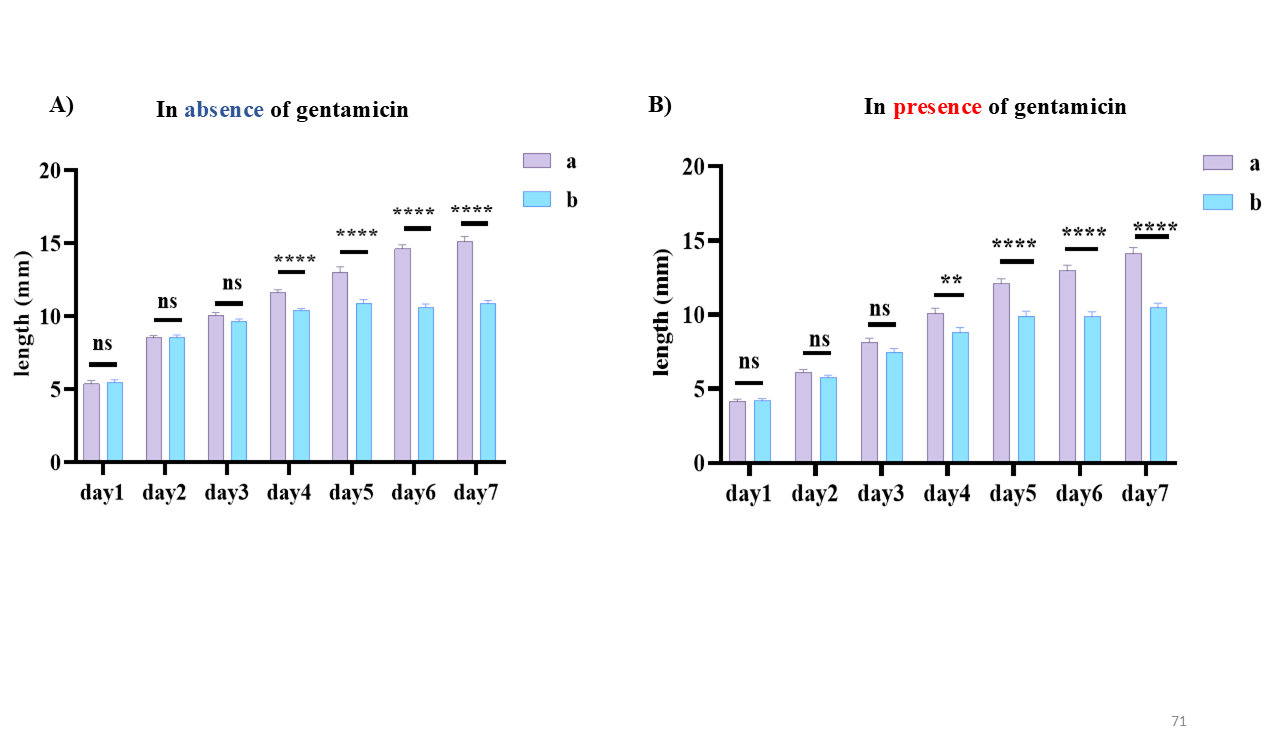

**Figure S7.** Length from the colony center towards the a) non-interacting and b) interacting zones of sibling colonies in A) the absence of gentamicin and B) the presence of gentamicin. Data presented as the mean ± SEM for N=3, where * is *P* ≤ 0.05, ** is *P* ≤ 0.01, *** is *P* ≤ 0.001, **** is *P* ≤ 0.0001, and ns signifies non-significant.

**Figure S8**

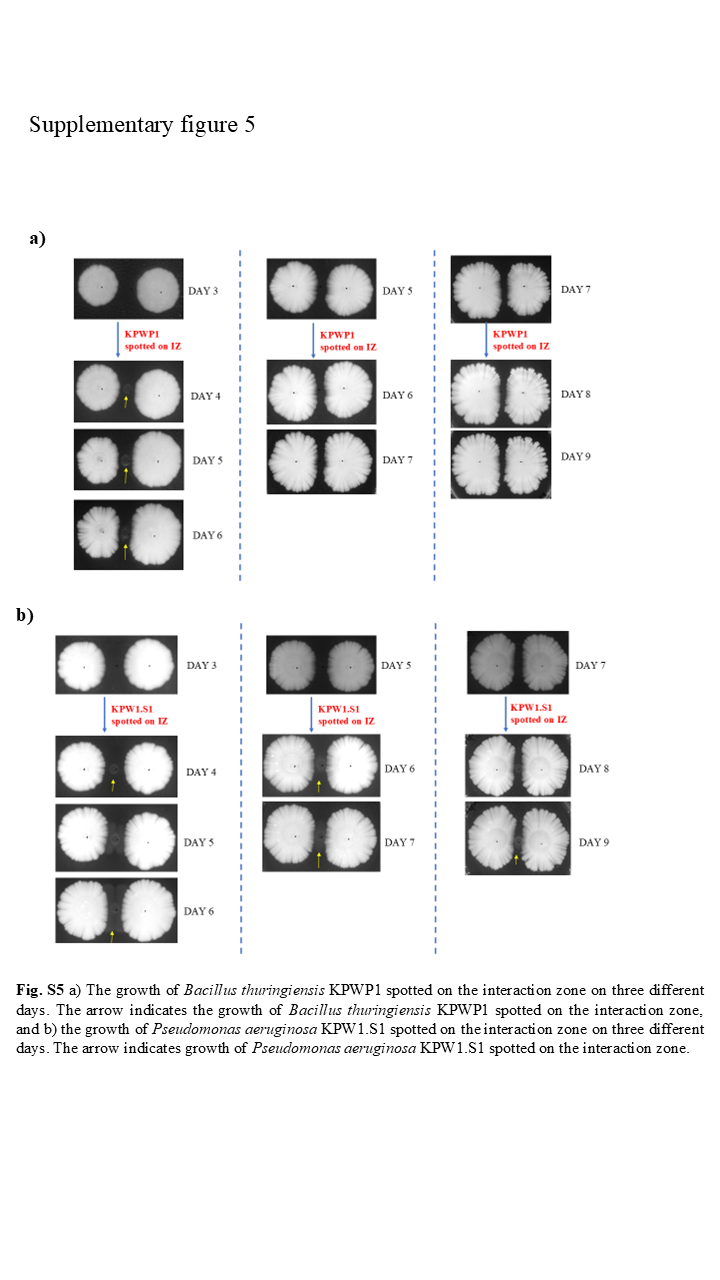

**Figure S8.** a) The growth of *Bacillus thuringiensis* KPWP1 spotted on the interaction zone on three different days. The arrow indicates the growth of *Bacillus thuringiensis* KPWP1 spotted on the interaction zone, and b) the growth of *Pseudomonas aeruginosa* KPW1.S1 spotted on the interaction zone on three different days. The arrow indicates growth of *Pseudomonas aeruginosa* KPW1.S1 spotted on the interaction zone.

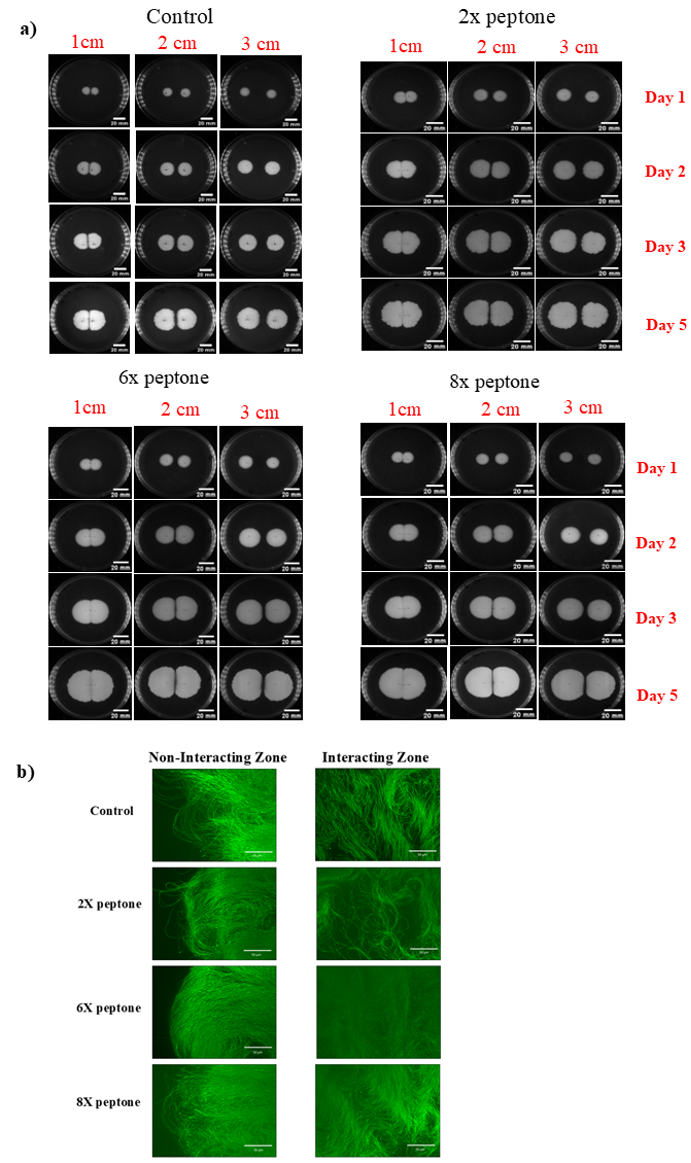
**Figure S9**

**Figure S9.**  a) interaction between sibling colonies of *Bacillus cereus* MSM-S1 in the presence of increased peptone concentrations, and b) arrangement of cells from the non-interacting and interacting zones during interaction between sibling colonies of *Bacillus cereus* MSM-S1 growing at a distance of 1cm on nutrient agar plates with 0.6% agar with increased peptone concentrations.
